# Capturing gait parameters during asymmetric overground walking using ultra-wideband radars: A preliminary study

**DOI:** 10.1101/2024.07.01.601550

**Authors:** Charalambos Hadjipanayi, Maowen Yin, Timothy G. Constandinou

## Abstract

This study investigates the deployability of commercially available impulse-radio ultra-wideband radar (UWB) sensors, in accurately detecting and analysing gait patterns during asymmetric overground locomotion. An adjustable knee brace was fitted on the right knee of 10 able-bodied participants, in five different confinement angles, during a 6-meter walking task to simulate asymmetric walking patterns. A computationally efficient signal processing framework extracts seven spatiotemporal gait parameters from six UWB radar signals, based on their joint Range-Doppler-Time representation. Gait asymmetry was quantified using primarily the symmetry ratio metric of step times. Validated against the gold standard motion capture system, radar-based gait parameters were estimated with 88.2-98.8% accuracy for all settings. By capturing step time symmetry ratios with 95.6±2.8% accuracy, the radar system can effectively distinguish between different gait impairment levels.

## I. INTRODUCTION

Disruption of gait pattern coordination, often resulting from musculoskeletal or neurological disorders, has a detrimental impact on an individual’s mobility, independence, health, and overall quality of life. Asymmetric gait patterns, which are particularly prevalent among the elderly population [1], can compromise physical function and increase the risk of falls [2], leading to potentially major injuries. Long-term monitoring of inter-limb gait asymmetries is thus critical for clinicians for tracking disease or injury progression, quantifying fall risk and assessing intervention or medication efficacy.

Many lab-based technologies have been developed for analysing gait patterns, however, research now focuses on home-oriented solutions, as they can potentially provide continuous, convenient and cost-effective patient monitoring. Existing home-based gait analysis systems, are commonly hindered by high cost, low portability, privacy concerns, user noncompliance and discomfort, making them inconvenient for longitudinal monitoring of elderly people. The latest advances in ultra-wideband (UWB) radar-on-chip technologies provide a safe, unobtrusive, affordable, portable, and low-power monitoring solution.

UWB radar sensors transmit high-bandwidth pulses towards a target of interest and based on time-of-flight measurements and backscattered signal modulation, information about the target’s motion can be recovered. These systems have been widely used in research for detecting vital signs and behavioural signals due to their high spatial resolution, and thus have the potential to detect small variations and asymmetries from gait patterns. Various studies have employed radar sensors for extracting spatiotemporal gait parameters based on the Doppler frequency information captured in the received radar signals [3], [4], [5]. Moreover, the study by Seifert et al. [6] explored the use of radars for capturing gait parameters from artificially-induced asymmetric treadmill walking using an adjustable knee brace and reported the accuracy of radar-based features for different knee brace settings. Treadmill walking, however, may influence an individual’s natural walking pattern, as it imposes invariance to gait parameters [7] and does not closely resemble realistic walking conditions in home environments.

Within this context, our study investigates the novel application of UWB radars for accurate detection and analysis of gait asymmetries during overground walking trials, expanding upon the treadmill-based study conducted by Seifert et al. [6]. In contrast to the methodology employed in their work, which focused on tracking Doppler frequency components solely over time using Doppler spectrogram analysis, our study adopts the joint range-Doppler-time (RDT) representation [5], [8]. This representation captures both temporal and spatial Doppler information making it ideal for overground walking applications. Using experimental data from 10 healthy individuals recorded during a 6-meter walking task, this work assesses the radar’s capability to capture gait parameters and quantify gait symmetry/asymmetry under a) normal and b) impaired walking, using an adjustable knee brace that restricts flexion of one knee. The accuracy of the extracted radar-based metrics is validated against marker-based motion capture (MOCAP) data, which currently serves as the gold standard for gait assessment.

## II. Methodology

### A. Experimental Setup

The experimental setup used for data capturing is illustrated in Fig. 1A. Radar data was captured using six independent impulse-radio ultra-wideband (IR-UWB) radar systems on a chip (XeThru X4M03, Novelda AS, Norway). All sensors were operating at a carrier frequency of 7.29GHz with a frame rate of 500Hz, allowing target radial velocities up to 5.15m/s to be unambiguously detected.

**Fig. 1.**
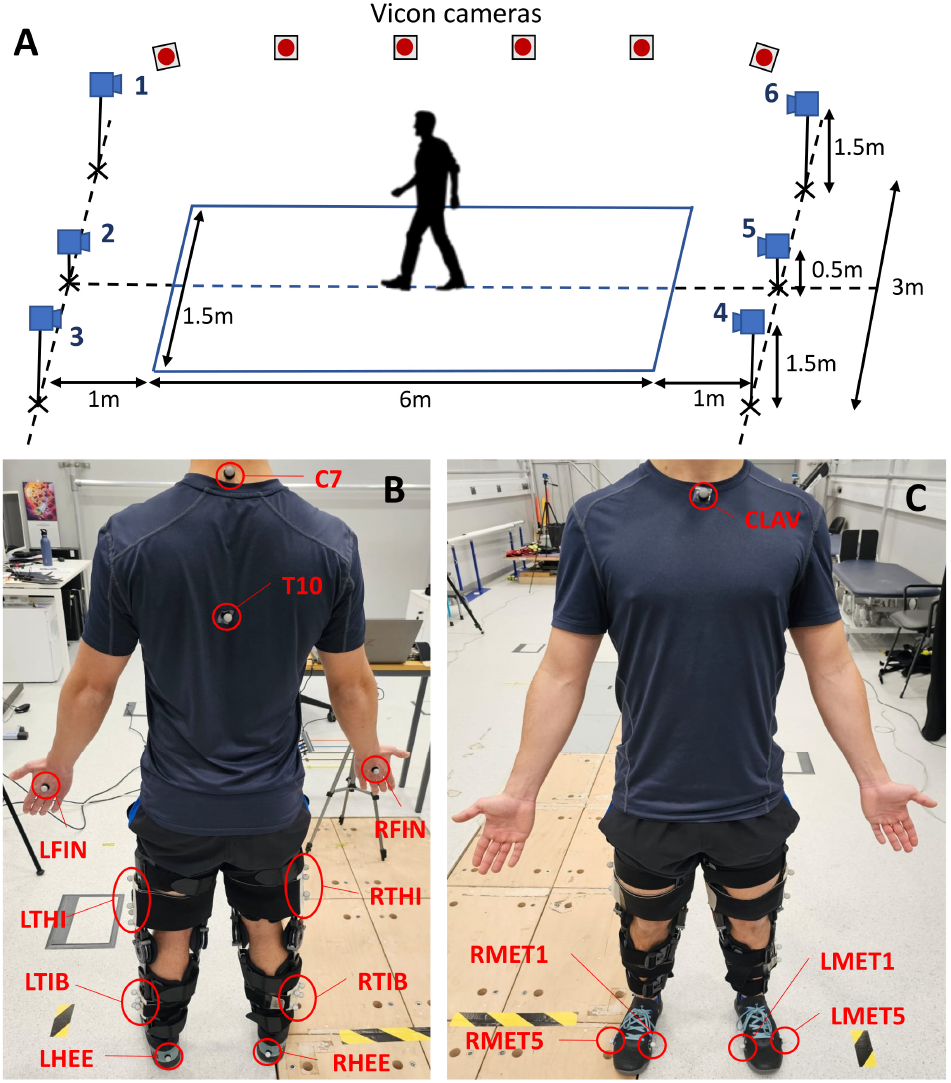
Experimental set-up employed in this study: (A) Location of radar sensors, (B&C) Placement of the 27 reflective markers on the ventral and dorsal parts of the subject’s body.

The kinematic data of body parts, serving as ground truth, were collected using a Vicon motion capture (MOCAP) system consisting of 28 high-precision infrared cameras (16 Vantage v8 and 12 Vero v2.2, Vicon Motion Systems, UK). A total of 27 passive reflective markers were attached to the participant’s body at landmarks shown in Fig. 1B and 1C.

Asymmetric gait patterns were simulated using an adjustable hinged knee brace (X-ROM, DJO Global, USA) fitted on the right leg, to restrict knee flexion. Since the presence of the brace affects the radar cross section (RCS) of the leg, a second identical knee brace was fitted on the left leg in an unrestricted configuration.

### B. Participant Information and Protocol

Ten healthy participants (6 males and 4 females, age: 26.5±2.8 years, height: 1.71±0.1 m, weight: 67.4±10.6 kg) were recruited to perform the 6-meter walking trials. The experimental trials, approved by the Imperial College London ethics committee (SETREC approval ref: 21IC6761), were conducted at the Biodynamics Laboratory (MSk Lab) at Imperial College London, UK. Participants were instructed to walk at their normal speed along the designated pathway for eight times, under six different right knee flexion confinement angles. Apart from free unrestricted motion, the knee flexion confinement angles set by the adjustable brace were 0°, 10°, 20°, 30°and 40°.

### C. Radar Signal Model

The transmitted radio frequency (RF) pulse of the coherent X4M03 IR-UWB sensor [9] can be described as

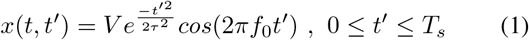

where *T*_*s*_ is the pulse repetition interval, *t*^*′*^ denotes fast time sampled at intervals of *T*_*f*_, where *T*_*f*_ *≪ T*_*s*_, *t* = *mT*_*s*_ is slow time, *m* is the pulse index, *V* is the peak amplitude, *f*_0_ is the carrier frequency and *τ* determines the -10dB pulse bandwidth. The received baseband radar signal from *K* body parts during motion, after the quadrature demodulation process, is approximated as

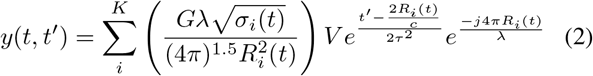

where *R*_*i*_ and *σ*_*i*_ are the radial displacement (range) and RCS of i^th^ body part, *c* is the speed of light, *λ* is the signal wavelength, and *G* is the antenna gain. By computing the short-time Fourier Transform (STFT) of the received signal across the slow-time dimension, the spectrum magnitude for the i^th^ body part is maximum at the frequency *f*_*i,p*_, given by

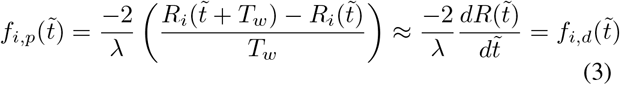

where *T*_*w*_ is the STFT window length, 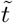 the new resulting slow-time variable and 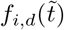 is the i^th^ body part’s instantaneous

Doppler frequency, assuming that the body part’s radial velocity is constant within interval *T*_*w*_. This shows that faster-moving body parts (e.g. feet) during walking will have peak STFT magnitudes at higher frequencies than slower-moving parts (e.g. torso). Additionally, the STFT magnitude for a body part is directly proportional to its RCS value, thus larger body parts produce stronger STFT components.

### D. Motion Capture Kinematic Data Analysis

Noise artefacts from the marker kinematic data are suppressed using a 4^th^ order Butterworth low-pass filter with cutoff frequency at 7Hz. The Foot-Velocity algorithm [10] is then used to detect heel-strike (HS) events, i.e the time instants of initial heel-ground contact during walking, from left and right foot marker trajectories.

### E. Radar Data Processing

The radar signal processing steps implemented in this study, based primarily on the methodology proposed in our previous work [5], are summarised in Fig. 2A.

**Fig. 2.**
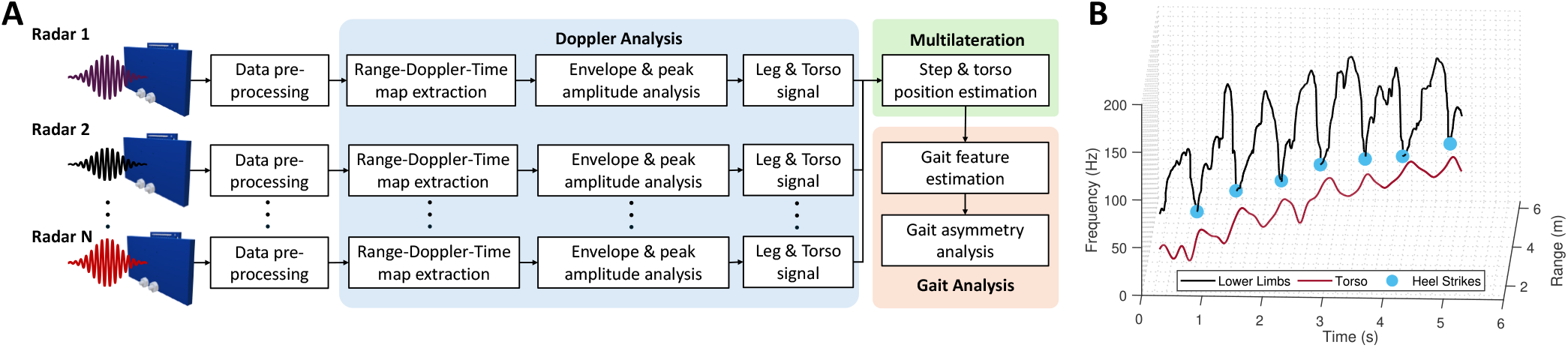
(A) Flowchart of radar signal processing approach. For this study, the total number of radars (N) is 6. A minimum of 3 radars is required for this approach to work. (B) Detected HS events from the extracted feet signal of a single radar.

#### Pre-processing

A 2^nd^ order Butterworth high-pass filter, with cut-off frequency at 2.4Hz, is used to remove radar returns from clutter with radial velocity below 0.05m/s. Noise artefacts are suppressed using the matched filtering method, where the transmitted signal in (1) is used as a template. Finally, a quadrature demodulation step is performed to convert the received RF signal into its baseband equivalent.

#### Doppler Analysis

The joint RDT map of the received baseband signal is extracted by computing its slow-time STFT

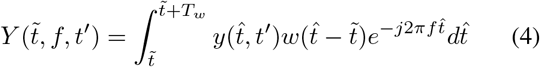

where *w* is the STFT window function of length *T*_*w*_ and *f* is the Doppler frequency dimension. In this study a 0.2s-long Kaiser window was used, with a roll-off coefficient of 10 and 95% overlap. Weaker Doppler components due to feet are enhanced using the motion enhancement approach proposed in [11]. The envelope signals of each RDT frame are extracted, similar to the approach followed in [6], using the percentile method [12]. The feet (or lower-limb) signal, consisting mainly of components from both feet, is extracted by tracking the envelope frequency extrema in both range and time, while the torso signal is captured by tracking the frequency value corresponding to the highest magnitude in each RDT frame. HS events are detected when the feet signal reaches a local minimum, as shown in Fig. 2B.

#### Multilateration - Data Fusion Stage

The torso and feet signals from each radar are combined using the weighted least squares multilateration approach [13] to estimate their three-dimensional (3D) position, accounting for random errors in range estimations. The estimated values of HS times from all radars are also integrated through a weighted average approach. Based on these unified values, the 3D position of feet during HS events is obtained. The weights assigned to each sensor in both methods are equal to the reciprocal of the estimated torso’s range by that radar. This accounts for signal quality degradation caused by path loss effects.

### F. Gait Parameter Extraction

The spatiotemporal relationship between consecutive HS is used to estimate gait parameters of high clinical importance from both systems, including:

- Step/stride time: Time intervals between consecutive HS times of opposite feet (step) or same foot (stride).
- Cadence: Reciprocal of the step time.
- Step/stride length: Distance covered by feet along the direction of travel between consecutive HS events of opposite feet (step) or same foot (stride).
- Walk ratio: Ratio of step length and cadence.
- Walking speed: Derivative of the torso’s position along the direction of travel.

The spatial features under consideration are exclusively determined based on displacements along the walking direction, due to larger multilateration errors of the radar system along the other directions. The estimated displacements along the medial-lateral direction serve solely to distinguish which HS events occur in the left or right foot.

The symmetry ratio is used in this study to quantify gait asymmetries, which is a widely adopted metric due to its straightforward interpretation [14]. The symmetry ratio is simply defined as the ratio of gait parameter values between the confined (right) and unconfined (left) leg. Step time and step length parameters are commonly used for symmetry ratio computation [14].

## III. Results AND Discussion

### A. Gait Parameters and Symmetry Ratio

A comparison of the estimated average radar-based parameters and their ground truth MOCAP values, over all participants, for all knee restriction angles is provided in Fig. 3. The accuracy of the radar system is indicated in the boxplots in Fig. 3, which is quantified using the mean absolute percentage error. The obtained results indicate that the radar system can accurately capture all considered gait features and their associated variability. Step/stride time, cadence, stride length, and walking speed are captured with more than 95.3% mean accuracy, while step length and walk ratio achieve at least 90.5% mean accuracy. The findings also suggest that with decreasing knee confinement (greater knee flexibility), there is an overall increasing trend in cadence, step/stride lengths, and walking speed, and a decreasing trend in step/stride times. No clear trend can be identified for walk ratios. Furthermore, there is no clear significant difference in the accuracy of extracted parameters across knee restriction angles, as evidenced by the highly overlapping boxplots.

**Fig. 3.**
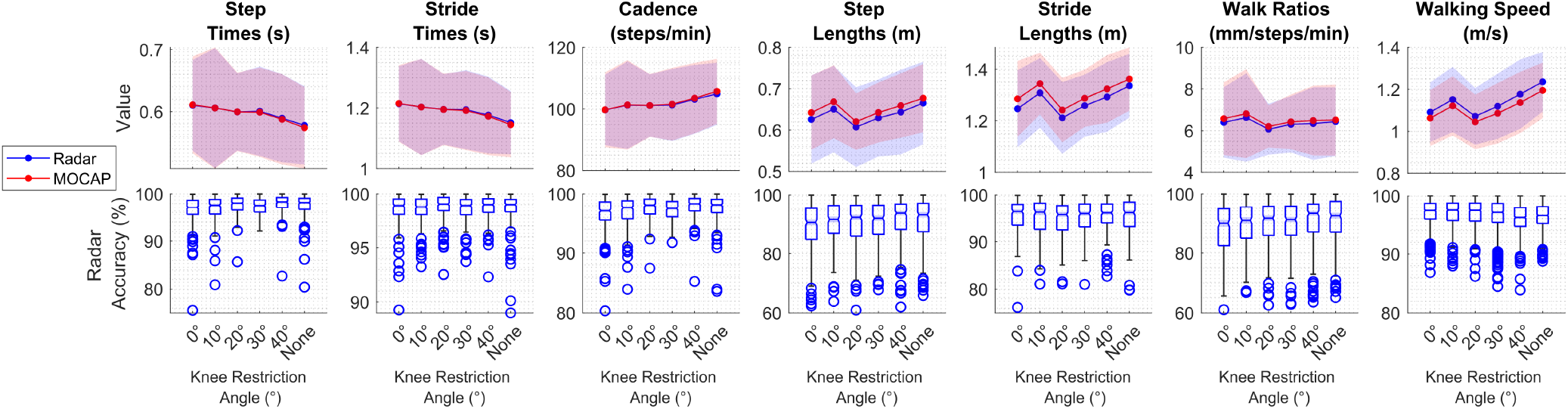
(Top) Average gait features extracted using radar and MOCAP systems for each knee confinement angle. Shaded areas correspond to one standard deviation from the mean value. (Bottom) Accuracy of radar-based gait parameters for each knee confinement angle, with blue circles denoting outliers.

Fig. 4 presents the symmetry ratio values for both step time and step length parameters. The step length symmetry ratio values for the MOCAP system (Fig. 4A) suggest no significant differences among different knee confinement angles. Although mean step length generally increases with larger knee confinement angles, left and right leg step lengths remain relatively constant. However, Fig. 4C reveals a more pronounced decreasing trend in step time ratio values with increasing knee confinement angles in the MOCAP system. Considering this, and given the radar system’s higher accuracy in detecting step times, only radar-based step time symmetry ratios are analysed (Fig. 4B). The radar system captures step time symmetry ratios with 95.6*±*2.8% accuracy, demonstrating its ability to detect the same decreasing trend as the MOCAP system. Finally, the correlation plots of the extracted step time symmetry ratios are shown in Fig. 5, comparing 0°and 20°knee confinement angles against no confinement. The results indicate that the radar system effectively distinguishes between cases of no and maximum knee confinement. In the 20°case, there is a higher degree of overlap between the two clusters, yet they can still be separated.

**Fig. 4.**
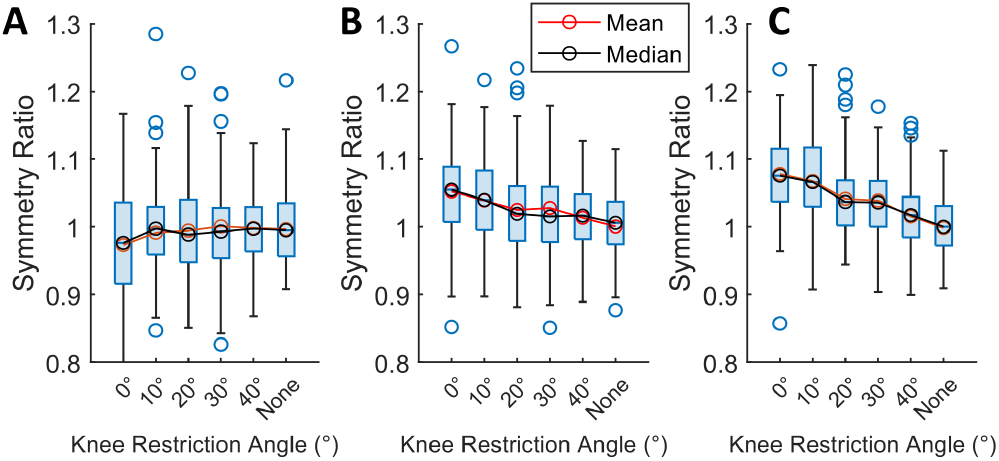
(A) Symmetry Ratio for step lengths using MOCAP. (B) Symmetry Ratio for step times using radar. (C) Symmetry Ratio for step times using MOCAP.

**Fig. 5.**
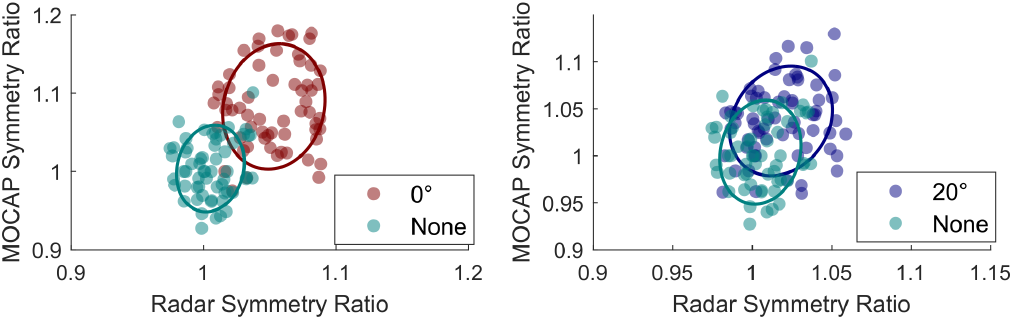
Correlation plots for step times from both systems, comparing (Left) no confinement with 0°and (Right) no confinement with 20°. Only step time values within the interquartile range are shown.

### B. Study Limitations

All experimental trials were conducted in a controlled lab-based environment where participants were walking purely in one direction in each walking segment, which differs from real in-home conditions. A larger cohort, including patients with gait impairments, is necessary for a more comprehensive assessment of radar capabilities. Additionally, gait parameters related to gait cycle phases, e.g. swing/stance time [14], should be also considered for gait asymmetry calculation. Although six radar sensors were employed to improve gait analysis accuracy, this may be impractical for home applications. Finally, radar placement must be optimised to minimise multilateration errors, allowing for the extraction of medial-lateral features, such as step width.

## IV. CONCLUSIONS

This study demonstrated that UWB radar sensing can provide sufficient information for assessing symmetric as well as asymmetric human overground walking. As a completely unobtrusive and low-cost solution, radar sensing is a promising alternative to lab-based systems, for continuous and long-term gait monitoring of people with musculoskeletal or neurological disorders at their own homes.

